# Aging-associated dysbiosis increases susceptibility to enteric viral infection in *Drosophila*

**DOI:** 10.1101/156455

**Authors:** Christine Sansone, Jonathan Cohen, Beth Gold, Wenhan Zhu, Ana M. Misic, Daniel P. Beiting, Sebastian E. Winter, Sara Cherry

## Abstract

Age is associated with increased susceptibility to enteric infections, but the molecular mechanisms are unclear. We find that aged *Drosophila* are more susceptible to enteric viral infections and that this increase in susceptibility is due to the aged microbiota, since depletion of the microbiota or reconstitution with a young microbiome suppressed infection. Metagenomic analysis of the aged microbiome revealed dysbiosis with an increased abundance in reactive oxygen species (ROS) producing pathways. This aged microbiota drives intestinal ROS production and we could restore immune function in old flies by reducing ROS genetically or pharmacologically. Moreover, we found that reconstitution of old flies with a cocktail of commensals, including *L. fructivorans* and heat-killed *A. pomorum*, could fully restore immunity. Altogether, these findings provide a mechanistic link between age-dependent dysbiosis and antiviral immunity and show that we can restore innate protection in aged animals, suggesting that this is a treatable and reversible state.

## Introduction

Aging is associated with progressive decline in physiological functions, including dysfunction of the immune system, which results in increased susceptibility to a variety of infections (Lopez-Otin et al., 2013; Ponnappan and Ponnappan, 2011). In particular, in the intestine, gastroenteritis is common in the elderly and is caused by a variety of pathogens including *Salmonella*, *Shigella*, *E. coli*, norovirus, rotavirus, astrovirus, and adenovirus (Jagai et al., 2014; Kirk et al., 2012; Kirk et al., 2010; Strausbaugh et al., 2003). Furthermore, aging animals have an altered microbiome, termed dysbiosis, which is implicated in diverse disease states from immune dysfunction to metabolic disorders and autoimmunity (Clemente et al., 2012; Hollister et al., 2014; Owyang and Wu, 2014). Many hallmarks of aging, including increased inflammatory signaling and oxidative stress, are due in part to the dysbiotic microbiota, since a healthy mileu can be restored in some contexts upon loss of the microbiome (Ayyaz and Jasper, 2013; Guo et al., 2014; Lakshminarayanan et al., 2014; Man et al., 2014). Despite the clear public health impact, the molecular links between aging, immunity, and the microbiome are only beginning to be defined.

From flies to humans, there are conserved physiological changes associated with age. In particular, aged flies have intestinal dysbiosis characterized by increases in the number of commensals as well as shifts in the community structure (Claesson et al., 2011; Clark et al., 2015; Guo et al., 2014; Li et al., 2016; Ren et al., 2007; Wong et al., 2011). In addition, aged flies manifest increased inflammatory signaling and the induction of reactive oxygen species (ROS) in the gut (Ayyaz and Jasper, 2013; Buchon et al., 2009; Buchon et al., 2014; Guo et al., 2014; Kim and Lee, 2014). The consequences of these changes on enteric viral infections have not been explored.

Enteroviruses are a widespread class of enteric picornaviruses that commonly infect humans and cause a range of clinical symptoms from asymptomatic to meningitis (Abzug, 2014; Jubelt and Lipton, 2014; Muehlenbachs et al., 2014). Moreover, aging leads to an increase in susceptibility to many enteric viral infections in humans (Jagai et al., 2014). The mechanisms at play are poorly understood. Enteroviruses also infect insects and the picorna-like virus of *Drosophila*, Drosophila C virus (DCV), is a widespread pathogenic enterovirus of fruit flies (Jousset, 1976). Arthropod-borne viruses (arboviruses) are another group of viruses of global importance. Infection of the insect vector occurs orally during the blood meal, while infection of vertebrate hosts is through an insect bite (Attardo et al., 2005; Hansen et al., 2014; Raikhel and Dhadialla, 1992). Viruses within this blood meal infect intestinal epithelial cells to establish infection, as is the case for many enteric infections in mammals (Davis and Engstrom, 2012; Steinert and Levashina, 2011; Weaver and Barrett, 2004). Moreover, there has been a resurgence of vector-borne viral pathogens, which have become an increasing source of worldwide morbidity and mortality in humans and livestock.

To establish infection, enteric viruses must overcome barrier immunity within the intestinal environment. It is clear that the microflora within the intestinal tract plays a fundamental role in gut immunity (Buchon et al., 2013; Charroux and Royet, 2012; Lee and Brey, 2013; Sommer and Backhed, 2013), and the microbiota can play diverse roles in enteric infection (Hegde et al., 2015; Karst, 2016; Sansone et al., 2015). In young flies, peptidoglycan of the Gram-negative commensal *Acetobacter pomorum* primes NFkB signaling in enterocytes to promote antiviral defense through the induction of the antiviral cytokine Pvf2. A second signal that requires sensing of viral infection by the enterocytes, promotes Pvf2 expression in the intestine. This cytokine binds to the receptor tyrosine kinase PVR, and activates an antiviral cascade in the intestinal epithelium (Sansone et al., 2015). Since particular components of the microbiota (e.g. Gram-negative commensals) were protective in young animals, we set out to explore how aging and the old microbiota impacts intestinal antiviral immunity.

We found that with age, *Drosophila* become more susceptible to diverse enteric viral infections. This is due at least in part to their inability to induce the antiviral cytokine Pvf2 upon viral infection. This shift in susceptibility to viral infection occurs in middle age and is transferrable; the young microbiota is protective, while the old microbiota was detrimental to young animals. Functional analysis of the microbiota indicated that the old microbiota harbored increased ROS pathways. We found that old flies display an increase in ROS in the intestine, and that by decreasing ROS either pharmacologically or genetically, we could suppress the age-dependent increase in infection and restore antiviral Pvf2 expression. This suggests that microbiota-dependent ROS was responsible for the increased susceptibility to infection. Since the young microbiota was protective, we examined whether individual commensals when monoassociated with old animals could restore antiviral defense. We found that the Gram-positive commensal *Lactobacillus fructivorans* could suppress age-dependent increase in infection, but not to the level of young conventional flies. This was due to the fact that this commensal lacked the DAP-type peptidoglycan required for Pvf2 induction. Therefore, we created gnotobiotic animals harboring *L. fructivorans* along with heat-killed *A. pomorum*, which could fully restore immunity and antiviral Pvf2 induction. Therefore, this probiotic cocktail could revert the immune function of aged animals to that of young healthy animals. Altogether, these findings mechanistically link aging-dependent dysbiosis with increased susceptibility to enteric infection and suggest that understanding these interactions can lead to new interventions.

## Results

### Middle-Aged Flies are More Susceptible to Enteric Viral Infection

Flies live approximately 60 days and physiological changes associated with age are accumulative (He and Jasper, 2014; Iliadi et al., 2012). 4-7 day old animals are young adults, while middle-aged animals are 32-35 days old. We orally challenged these middle-aged flies (32-35 days old), which we will refer to as old, with a panel of viruses that we previously found can orally infect flies (Sansone et al., 2015; Xu et al., 2013), including Drosophila C Virus (DCV), a natural enteric pathogen of flies, as well as Vesicular Stomatitis Virus (VSV), and Dengue Virus (DENV) which are arboviruses belonging to two disparate families (*Rhabdoviridae* and *Flaviviridae*, respectively). Older flies were more susceptible to enteric infection than young flies as measured by confocal microscopy (Figure 1A) and RT-qPCR (Figure 1C). As we previously observed, infection was largely restricted to enterocytes in the posterior midgut, (Figure S1A) (Sansone et al., 2015). Moreover, we found that aging had a major impact on survival. Old flies orally challenged with DCV succumbed to infection, converting a largely non-pathogenic infection into a lethal one (Figure 1B). Altogether our data indicate that with age, flies become more susceptible to enteric viral infections.

**Figure 1.**
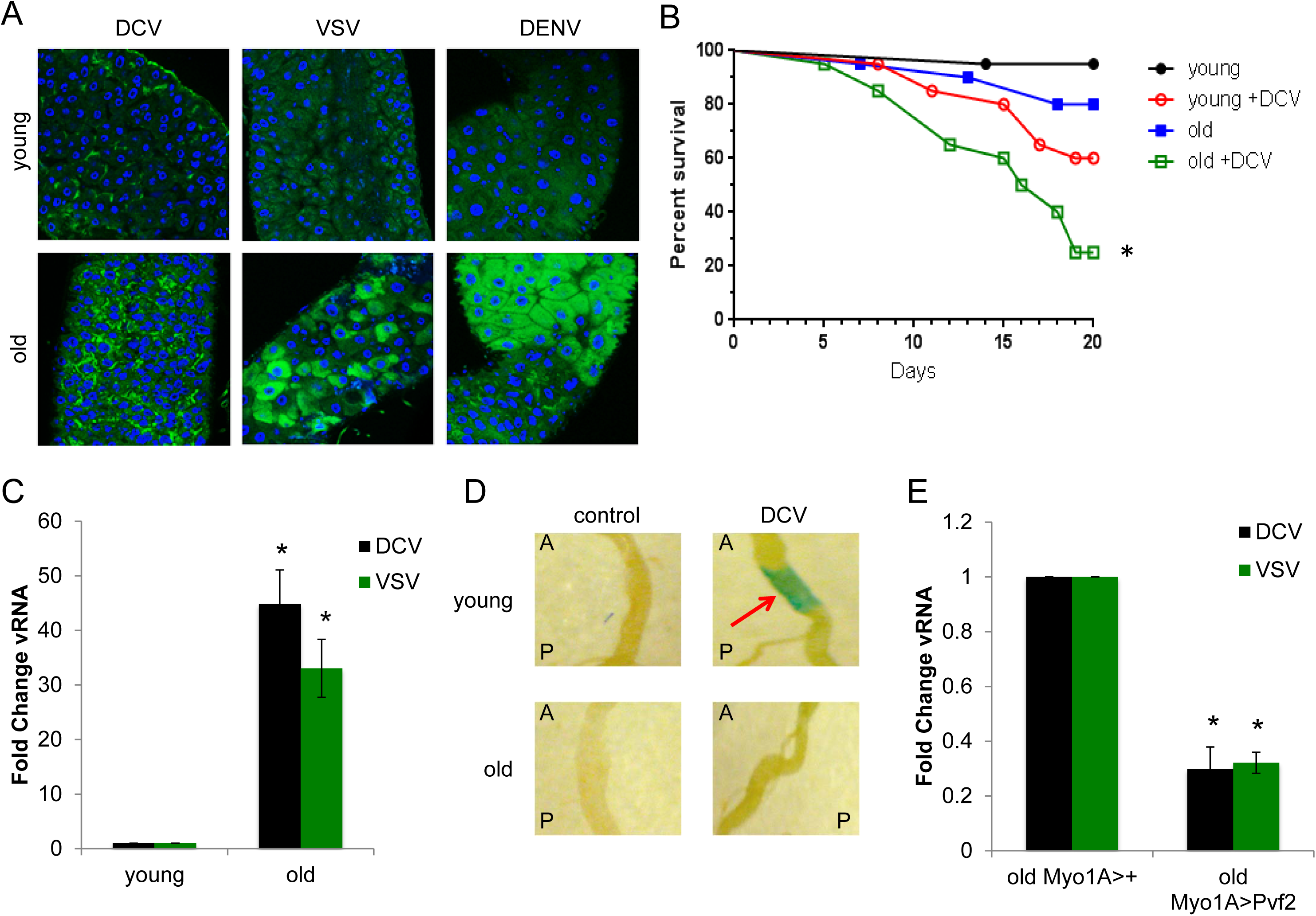
Older flies are more susceptible to enteric infection. (A) Representative confocal images of midguts from flies of the indicated age infected with the indicated viruses analyzed 7 dpi (DCV and VSV) or 10 dpi (DENV-2) (40x; virus-green, nuclei-blue) from n=3 experiments. (B) Flies of the indicated age were infected with the indicated viruses. Intestines were dissected and RT-qPCR analysis of viral RNA normalized to rp49 shown relative to control (young) 7 dpi with mean ± SD; n≥3; *p<0.05. (C) Percent survival of control (PBS-fed) or infected (DCV-fed) flies of the indicated age (n=3, p=0.0006, log-rank test). (D) Flies carrying a Pvf2 promoter-driven lacZ reporter (Pvf2-lacZ) of the indicated age were infected with vehicle or DCV and stained for beta-galactosidase activity at 3 dpi. A representative image of the posterior midgut and arrows indicate induction of lacZ expression (A, anterior; P, posterior). (E) Old flies of the indicated genotype were infected with DCV, intestines were dissected, and RT-qPCR analysis of viral RNA normalized to rp49 and shown relative to control 7 dpi with mean ± SD; n=3; *p<0.05.

### Older Flies Cannot Induce the Antiviral Cytokine Pvf2

Virus-induced cytokine production by enterocytes is essential for enteric antiviral protection in young animals (Sansone et al., 2015). Thus, we examined Pvf2 levels upon infection with age to determine if this was a cause of increased susceptibility to enteric viral infection. First, we monitored Pvf2 levels using transgenic flies that carry a lacZ reporter downstream of the endogenous Pvf2 promoter (Choi et al., 2008). As expected, young animals show an increase in lacZ activity in the posterior midgut upon DCV infection (Figure 1D). In contrast, older flies are unable to induce Pvf2 (Figure 1D). Second, we monitored the levels of Pvf2 mRNA by RT-qPCR and also observed no induction in older animals (Figure S1B). To determine if the lack of antiviral Pvf2 induction was driving the increased susceptibility to viral infection in older flies, we ectopically expressed Pvf2 in the intestinal epithelium of older flies. Indeed, ectopic expression of Pvf2 in older flies was protective, presenting with decreased viral infection (Figure 1E).

### The Aged Microbiota Increases Susceptibility to Enteric Infection

As they age, flies present with dysbiosis characterized by increased numbers and shifts in the community structure (Clark et al., 2015; Guo et al., 2014; Li et al., 2016; Ren et al., 2007; Wong et al., 2011). Since our animals were only middle-aged, we first explored the microbiome of our flies and found that the older flies presented with a ~2-log increase in bacteria per gut, as measure by colony forming units (Figure S2A), accompanied by increased inflammatory signaling in the intestine (Figure S2B). To determine whether the microbiota was playing a role in the increased susceptibility of older flies, we first ablated the microbiota with a cocktail of antibiotics we previously used (Sansone et al., 2015) and verified ablation (Figure S2C). Next, we orally challenged these flies with DCV or VSV. As previously shown, we found that the young microbiota is protective from enteric infection (Figure 2A) (Sansone et al., 2015). In contrast, the increased susceptibility of older flies was suppressed by antibiotic treatment (Figure 2A-B).

**Figure 2.**
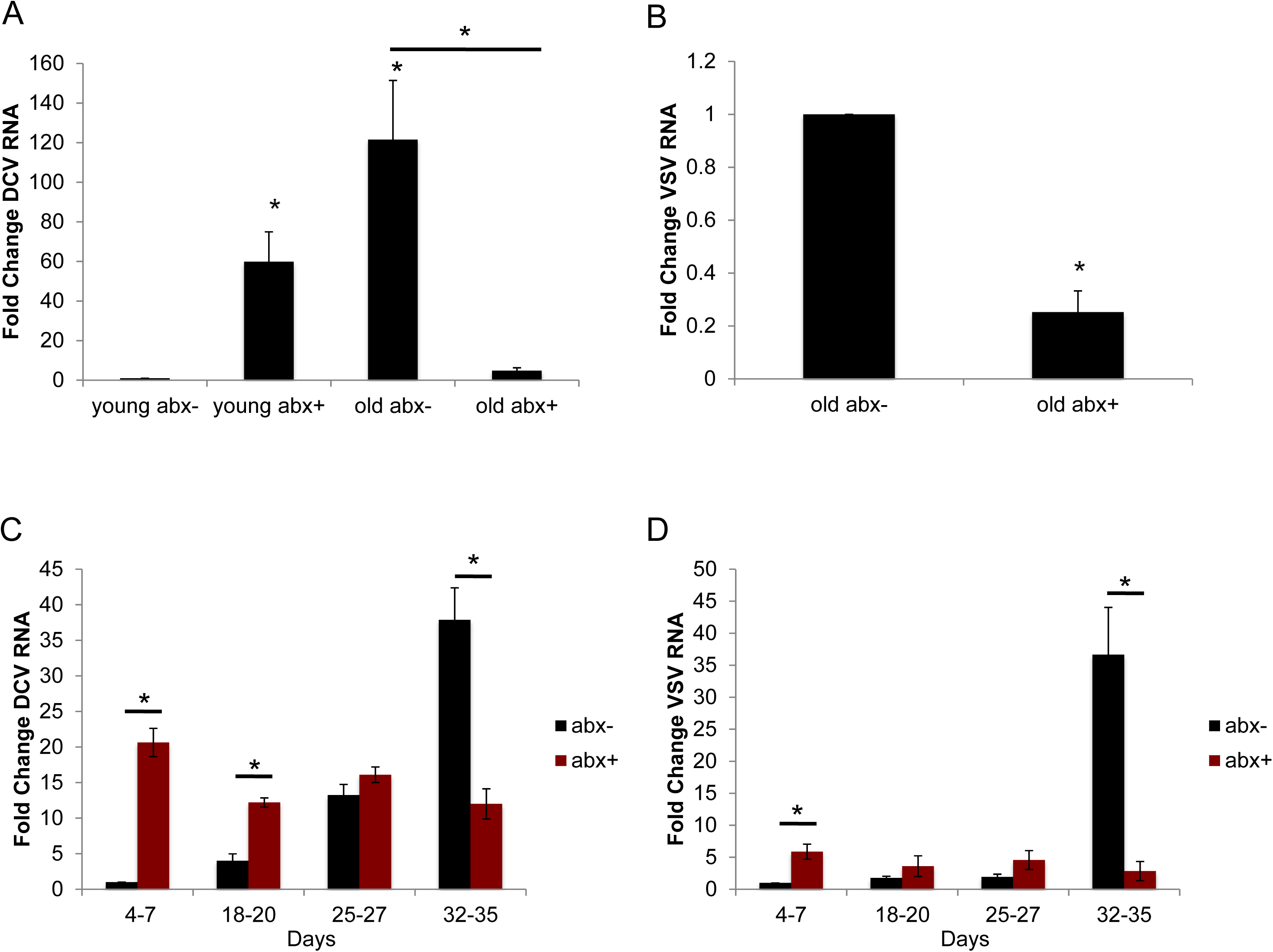
The microbiota of older *Drosophila* is detrimental to enteric antiviral defense. (A-D) Control or antibiotic treated flies of the indicated age were infected with (A,C) DCV or (B,D) VSV, intestines were dissected, and RT-qPCR analysis of viral RNA was normalized to rp49 and shown relative to control 7 dpi. Mean ± SD; n≥3; *p<0.05.

These data suggest that the microbiota of older animals is detrimental to antiviral defense. To determine the age at which the animals become more susceptible to infection and when the microbiota becomes harmful, we collected animals from different ages, antibiotic treated them, and then orally challenged with DCV or VSV. We found that over time, flies harbor increasing numbers of bacteria (Figure S2D). Further flies older than 27 days presented with increased bacterial counts and viral infection was suppressed with antibiotics (Figure 2C-D, S2D).

We set out to test if the microbiota is sufficient to drive changes in viral susceptibility and Pvf2 induction. For these studies, we performed fecal transfers between young and old flies. First, we ablated the microbiota of young or old flies with antibiotics and then we created gnotobiotic animals by feeding the flies the feces from either young (4-7 day old) or old (32-35 day) flies. These flies were then orally challenged with DCV. Young flies that had been reconstituted with the microbiota from older flies showed an increase in infection, bacterial populations (colony forming units), and inflammatory signaling, while the older flies reconstituted with the young microbiota were protected to the level of young animals (Figure 3A-C). In addition, these older flies reconstituted with the young microbiota were protected from viral infection and displayed decreased bacterial counts and inflammatory signaling compared to old flies harboring the microbiota from older flies (Figure 3A-C). These data demonstrate that the microbiota impacts susceptibility to infection and that the young microbiota is protective, while that of the older flies is detrimental to immune defense. Additionally, we tested whether older flies reconstituted with a young microbiota that restricted viral infection were now able to induce antiviral Pvf2 upon infection. Indeed, we observed wild-type levels of Pvf2 induction in these gnotobiotic virally infected animals (Figure 3D).

**Figure 3.**
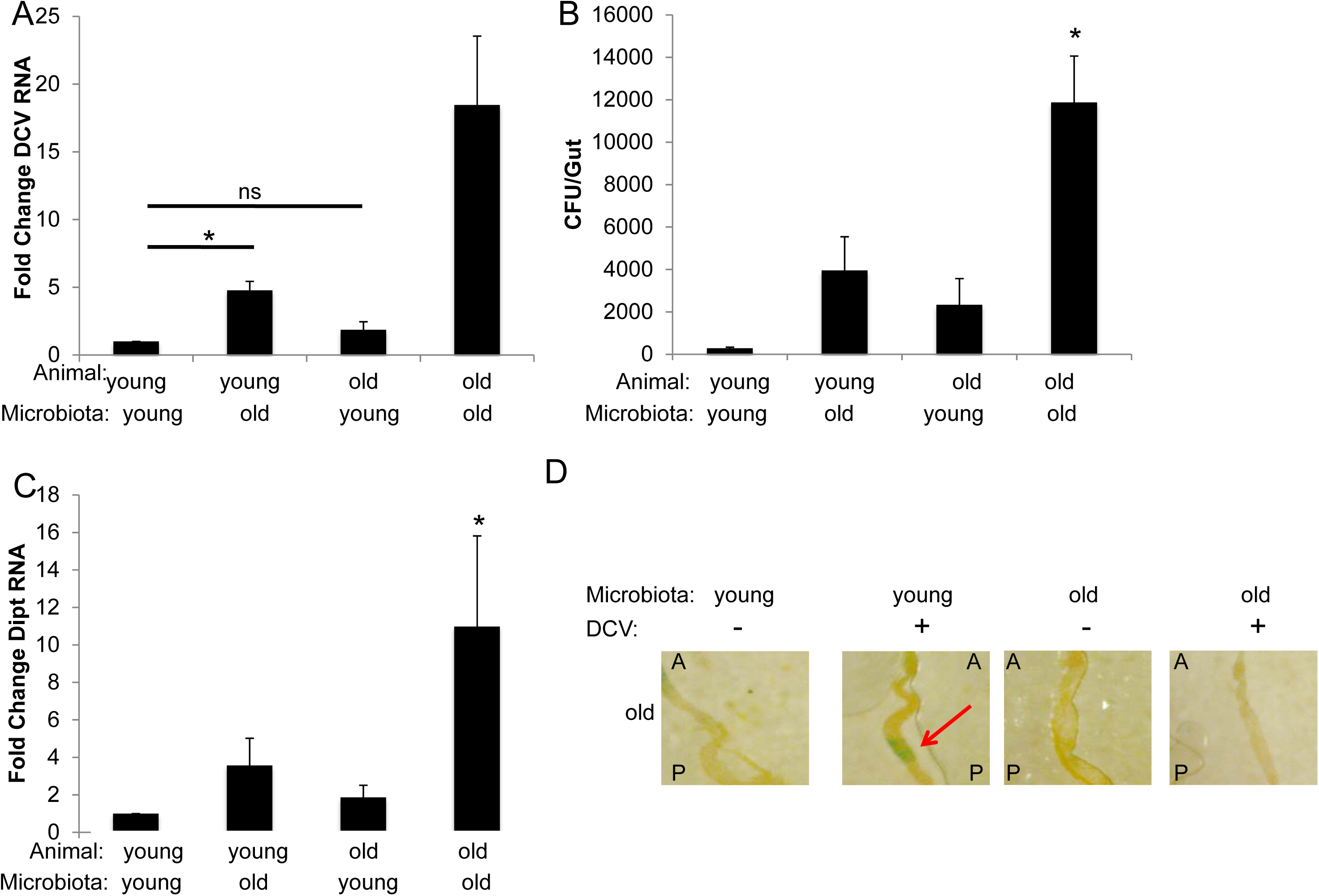
The young microbiota is protective. (A,C) Flies of the indicated age were associated with the indicated microbiota and infected with DCV. Intestines were dissected and RT-qPCR analysis of viral RNA (A) or diptericin RNA (C) was normalized to rp49 and shown relative to control 7 dpi. Mean ± SD; n=4; *p<0.05. (B) The number of colony forming units (CFU) from guts of flies of the indicated aged were associated with the indicated microbiota. Mean ± SD; n=4; *p<0.05. (D) Flies carrying a Pvf2 promoter-driven lacZ reporter (Pvf2-lacZ) of the indicated age were reconstituted with the indicated microbiota. Flies were then infected with vehicle or DCV and stained for betagalactosidase activity at 3 dpi. A representative image of the posterior midgut and arrows indicate induction of lacZ expression (A, anterior; P, posterior).

To elucidate how the composition of the fecal microbiome changes with age, we employed shotgun metagenomics on the feces of either young or old flies, and found that the microbiome of young animals is predominated by *Lactobacillus plantarum.* Indeed, Gram-positive bacteria make up 81% of the young microbiota (Figure 4A). In contrast, in old animals, the microbiome is predominated by two closely related Gram-negative commensals, *Acetobacter pomorum* and *Acetobacter pastuerians* (Figure 4A). Principle component analysis demonstrated that the predicted coding capacity of the microbiome in young flies and old flies form distinct states (Figure S3).

**Figure 4.**
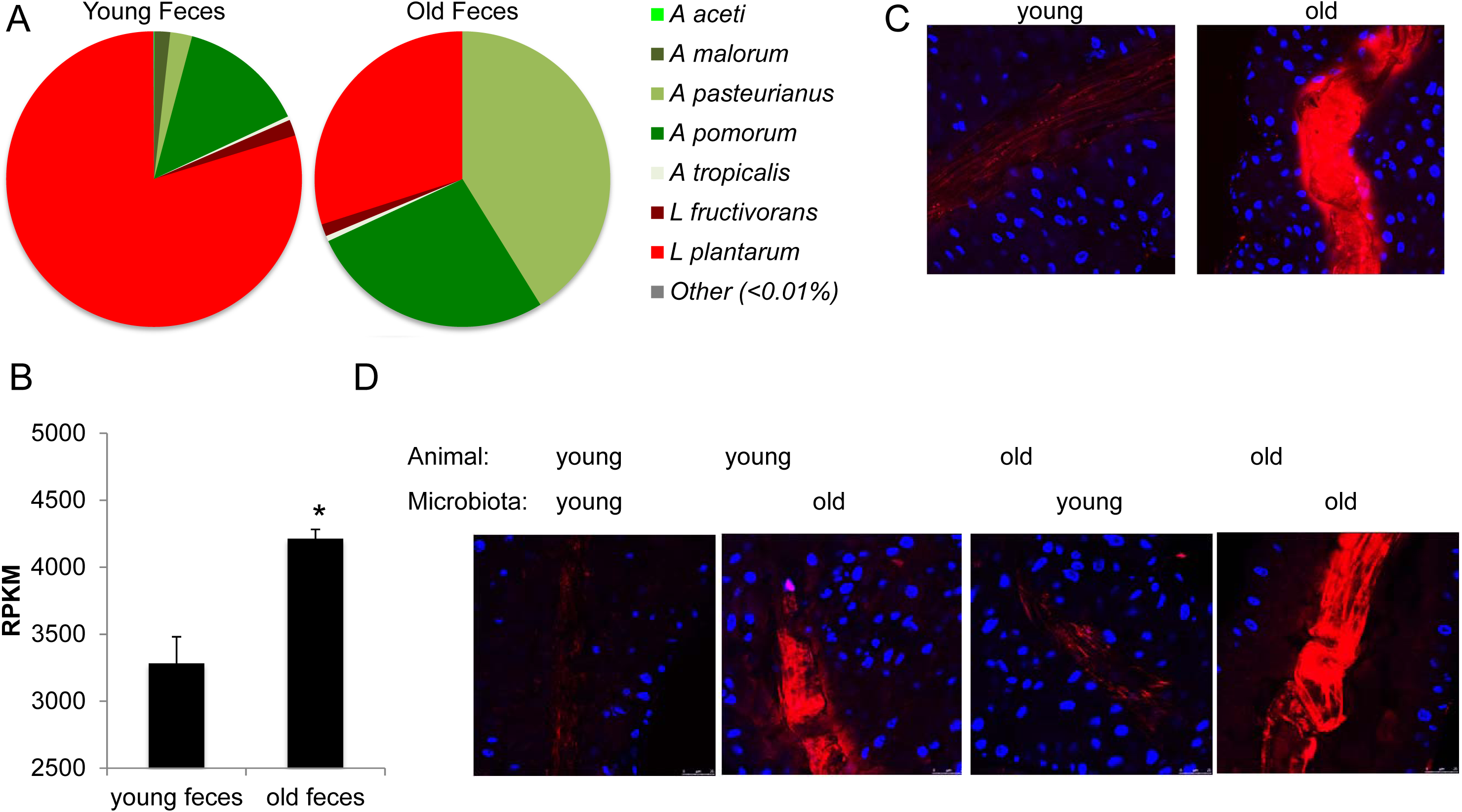
The old microbiota is associated with increased oxidative stress. (A,B) DNA extracted from the fecal contents of young and old flies was analyzed by shotgun metagenomic sequencing. (A) The relative abundance of bacterial species from indicated feces. Mean, n=2. (B) Reads mapped to ROS pathways in the KEGG orthology database were tallied from the indicated feces. (C,D) Representative confocal image of midguts of the indicated age (C) or associated with the indicated microbiota (D) and fed ROSStar 550 to visualize oxidative stress. (40x; oxidative stress-red, nuclei-blue) from n=3 experiments.

To explore the changes in the function of these different microbial communities, we mapped our metagenomic reads to the KEGG orthology database to identify pathways that are altered with age. We found that oxidative stress and reactive oxygen pathways were significantly increased in old animals (~30%; Figure 4B). Since this shift in community structure is accompanied by a 100-fold increase in bacteria (Figure S2A), these data suggest a ~130-fold increase in ROS pathways in older flies. Previous studies have shown that the aging gut has high ROS levels (Buchon et al., 2009; Guo et al., 2014; Hochmuth et al., 2011). We used a previously established ROS sensitive dye to measure the levels of ROS in the aged gut (Luo et al., 2016). We found that there is a dramatic increase in ROS in the older fly intestine (Figure 4C). We next tested the role of the microbiota in driving ROS production. Young flies reconstituted with an old microbiota displayed an increase in ROS compared to young flies with a young microbiota (Figure 4D). Likewise, old flies reconstituted with a young microbiota displayed a decrease in ROS compared to old flies harboring an old microbiota (Figure 4D). These data show that the altered microbiota drives increased intestinal ROS.

### Aging-Dependent Oxidative Stress Increases Susceptibility to Infection

These results raise the possibility that high levels of ROS are detrimental to antiviral defense. To test this, we induced oxidative stress in young animals by feeding them paraquat, a potent inducer of ROS (Biteau et al., 2008; Bus and Gibson, 1984; Choi et al., 2008) (Figure 5A). We found that young paraquat fed animals have elevated level of intestinal ROS (Figure 5A) and were more susceptible to viral infection (Figure 5B). Furthermore, these young paraquat treated flies were unable to induce antiviral Pvf2 expression (Figure 5C), suggesting high levels of ROS are detrimental to antiviral defense.

**Figure 5.**
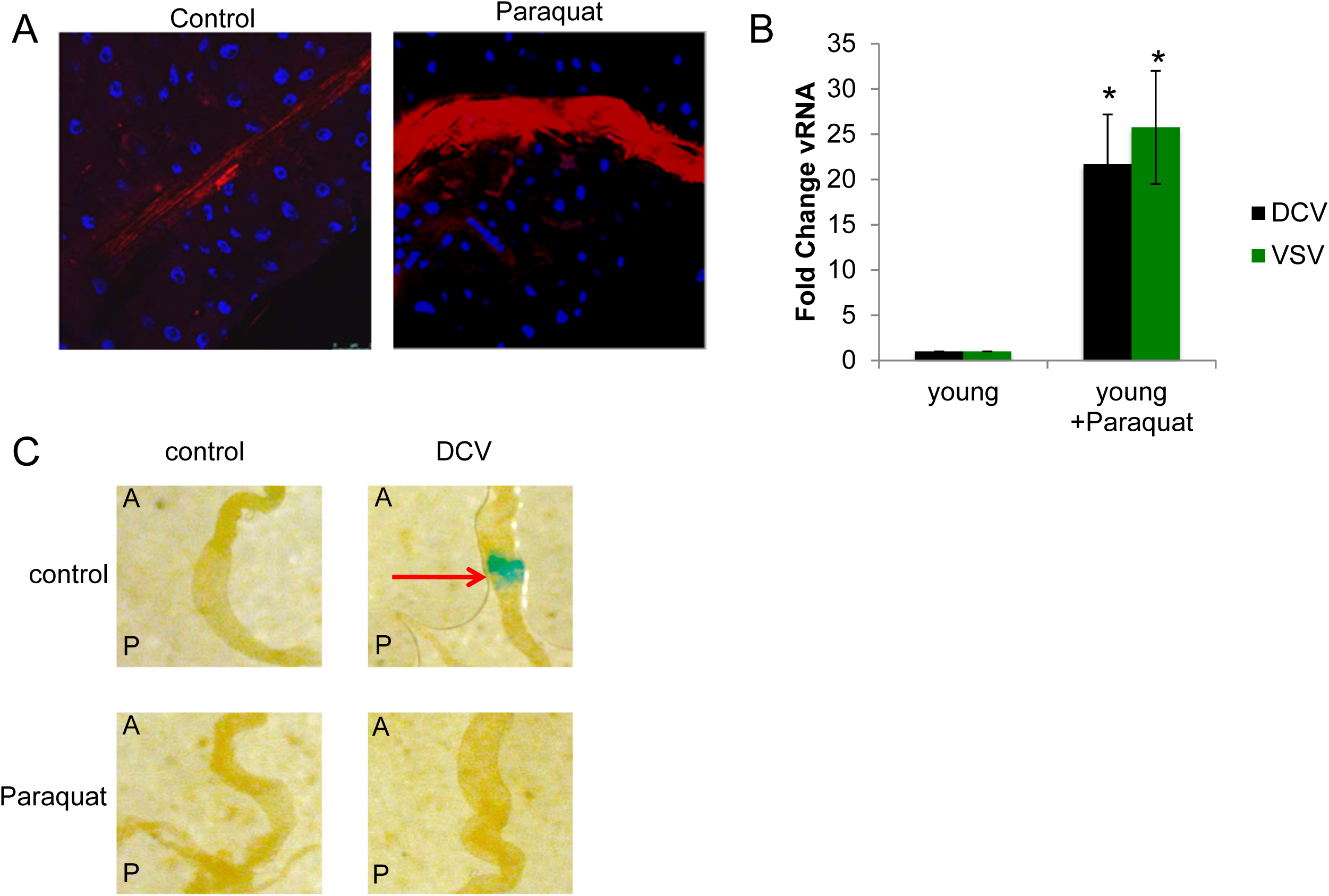
Inducing oxidative stress in young animals leads to increased susceptibility to enteric infection. (A) Confocal images of midguts from young flies fed either vehicle or paraquat (40x; oxidative stress-red, nuclei-blue). Representative images shown from n=3 experiments. (B) Young flies were paraquat treated and infected with the indicated viruses. Intestines were dissected and RT-qPCR analysis of viral RNA was normalized to rp49 and shown relative to control 7 dpi. Mean ± SD; n≥3; *p<0.05. (C) Young flies carrying a Pvf2 promoter-driven lacZ reporter (Pvf2-lacZ) were paraquat treated and then infected with vehicle or DCV and stained for beta-galactosidase activity at 3 dpi. A representative image of the posterior midgut and arrows indicate induction of lacZ expression (A, anterior; P, posterior).

Next, we reasoned that reducing ROS in older flies without altering the microbiota directly may induce protective immunity. Enterocytes produce ROS using two distinct NADPH oxidases, NOX and DUOX (Ayyaz and Jasper, 2013; Buchon et al., 2014; Kim and Lee, 2014). We first genetically altered NADPH oxidase levels in the intestine using previously validated *in vivo* RNAi to deplete either NOX or DUOX (Bae et al., 2010; Buchon et al., 2014; Ha et al., 2005a; Jones et al., 2013). We took advantage of a heat shock-inducible driver, where we allowed the animals to age normally, and then we depleted either NOX or DUOX in animals 3 days prior to oral challenge which had no impact on bacterial load (Figure S4A). While depleting either NADPH oxidase in young animals had no impact on infection, depletion in older flies resulted in a significant decrease in viral infection (Figure 6A,B). Next, we ectopically expressed iRC, an antioxidant catalase that reduces ROS (Ha et al., 2005b), and observed a significant decrease in infection in older flies with no impact in younger animals (Figure 6C).

**Figure 6.**
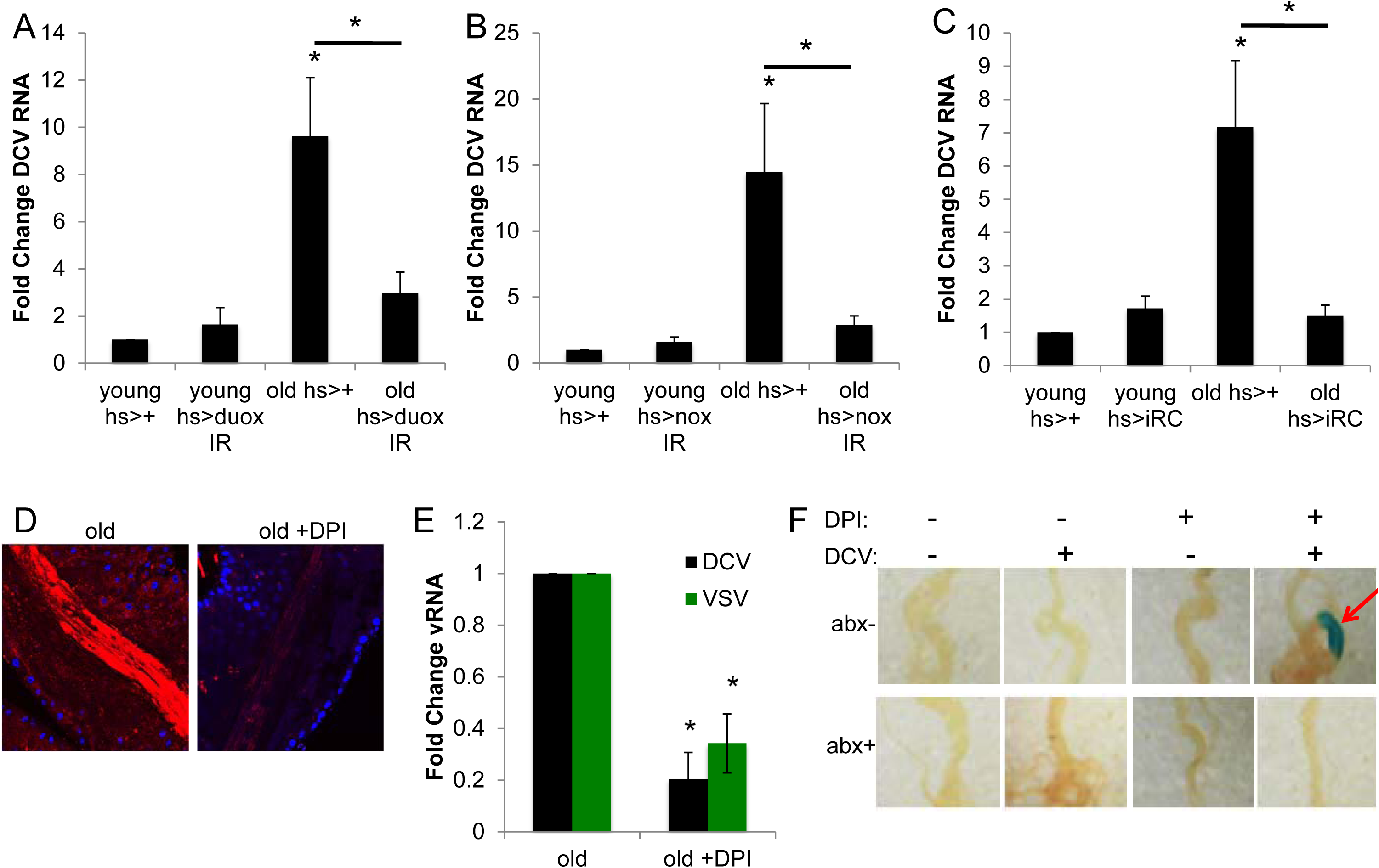
Oxidative stress in older *Drosophila* is detrimental to enteric infection. (A-C) Flies of the indicated genotype and age were infected with DCV. Intestines were dissected and RT-qPCR analysis of viral RNA was normalized to rp49 and shown relative to control 7 dpi. Mean ± SD; n≥3; *p<0.05. (D) Representative confocal image of midguts treated with either vehicle or DPI and fed ROSStar 550 to visualize oxidative stress. (40x; oxidative stress-red, nuclei-blue) from n=3 experiments. (E) Old flies were treated with the indicated drug and infected with the indicated virus. Intestines were dissected and RT-qPCR analysis of viral RNA was normalized to rp49 and shown relative to control 7 dpi. Mean ± SD; n≥3; *p<0.05. (F) Control or antibiotic treated old flies carrying a Pvf2 promoter-driven lacZ reporter (Pvf2-lacZ) were treated with the indicated drug, then infected with vehicle or DCV, and stained for beta-galactosidase activity at 3 dpi. A representative image of the posterior midgut and arrows indicate induction of lacZ expression (A, anterior; P, posterior).

In addition to genetic perturbations, we used a pharmacological inhibitor of NADPH oxidases, diphenyleneiodonium (DPI), which has been shown to decrease ROS (Cross and Jones, 1986; Ha et al., 2005a). Oral administration of DPI decreased ROS in old flies with no change in bacterial load (Figure 6D, S4B). Next, we orally challenged these flies with DCV or VSV and observed a significant decrease in viral infection compared to untreated flies (Figure 6E). Moreover, DPI treatment of older flies restored virus-induced Pvf2 expression (Figure 6F).

Our studies in young animals defined a two-signal requirement for antiviral Pvf2 induction: a signal from Gram-negative commensals to activate NF-kB and a second signal from viral sensing (Sansone et al., 2015). We found that increased ROS blocked Pvf2 induction (Figure 5C) and suppression of ROS restored virus-induced Pvf2 expression (Figure 6F). Since Gram-negative commensals are present in the old microbiota and NF-kB signaling is elevated (Figure S2B), these data suggested that ROS is blocking the virus-sensing signal. To test this model, we investigated whether DPI could restore Pvf2 expression in antibiotic treated animals. Indeed, we found that virus-induced Pvf2 expression was dependent on the microbiota (Figure 6F). Altogether, these data suggest that aging-dependent dysbiosis drives increases in ROS, which prevents viral sensing and thus antiviral cytokine production leading to increased susceptibility to enteric viral infection.

### A cocktail of two commensals restores immune function in old flies

These data suggest that aging prevents the virus-induced signal that drives Pvf2 expression. Since fecal transfers were sufficient to restore immunity (Figure 3A), we set out to test whether reconstitution of old flies with individual commensals could restore protective immunity. For these studies, we generated gnotobiotic animals harboring the two most prevalent Gram-positive (*L. plantarum* and *L. fructivorans*) or Gram-negative (*A. pomorum* and *A. pasteurians*) commensals that were identified in our metagenomics studies (Figure 4A). All four commensals efficiently colonized the aged gut (Figure S5A). Despite all four commensals suppressing the high inflammatory signaling present in the aged conventional animals (Figure S5B), only monoassociation with *L. fructivorans* significantly suppressed the age-dependent increase in susceptibility to enteric viral infection (Figure 7A, S5C). Monoassociation with the other commensals was not protective (Figure 7A). This is in direct contrast to young animals where we found that monoassociation with *A. pomorum* conferred protective immunity (Sansone et al., 2015).

**Figure 7.**
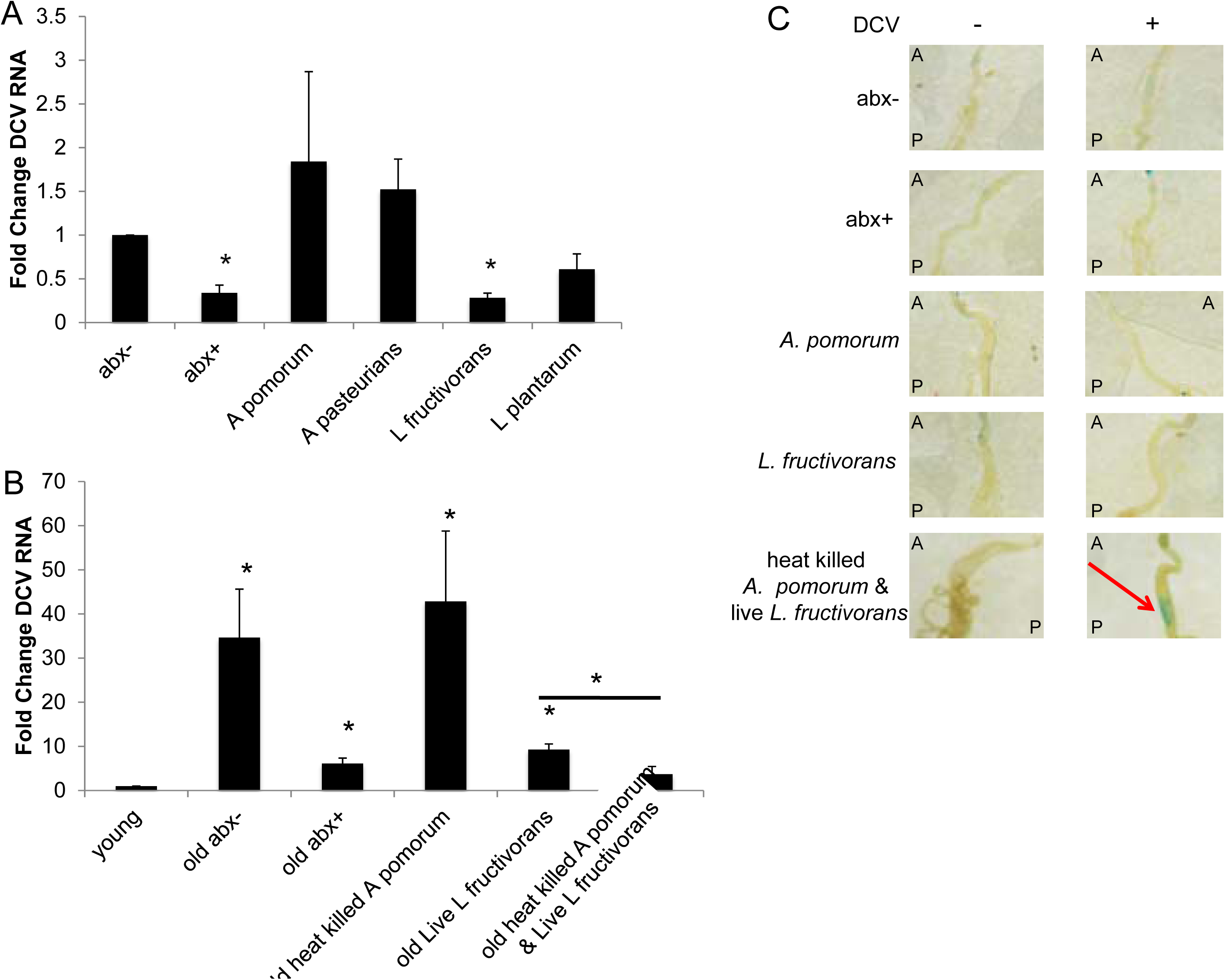
*L. fructivorans* and *A. pomorum* are required for antviral defense in older Drosophila. (A,B) Flies of the indicated age were associated with the indicated commensal and infected with DCV. Intestines were dissected and RT-qPCR analysis of DCV RNA was normalized to rp49 and shown relative to control 7 dpi. Mean ± SD; n≥4; *p<0.05. (C) Older flies carrying a Pvf2 promoter-driven lacZ reporter (Pvf2-lacZ) were associated with the indicated commensal and then infected with vehicle or DCV and stained for beta-galactosidase activity at 3 dpi. A representative image of the posterior midgut and arrows indicate induction of lacZ expression (A, anterior; P, posterior).

While the level of suppression by *L. fructivorans* was similar to that of antibiotic treated older animals, conventional young animals had lower levels of infection than either antibiotic treated or *L. fructivorans* gnotobiotic aged animals (Figure 7B). *Lactobacilli* spp. do not have DAP-type peptidoglycan and thus are unable to provide the priming signal for Pvf2 induction (Sansone et al., 2015). Indeed, monoassociatation with *L. fructivorans* was not sufficient to restore Pvf2 signaling (Figure 7C). We reasoned that the Gram-positive commensal was restoring the virus-induced signal without conferring the priming signal. Furthermore, we found that monocolonization with *A. pomorum* in old animals was neither protective nor sufficient to promote Pvf2 expression (Figure 7A,C). These data suggest that *A. pomorum* alone could not restore the virus-induced signal. Since we previously found that heat-killed *A. pomorum* is sufficient to prime NF-kB signaling for Pvf2 induction in young animals (Sansone et al., 2015), we reasoned that gnotobiotic animals harboring *L. fructivorans* along with heat-killed *A. pomorum* would provide both signals and restore complete immune function. Indeed, associating older flies with *L. fructivorans* and heat-killed *A. pomorum* was sufficient to suppress viral infection to the level of young conventional animals and restore Pvf2 expression (Figure 7B,C). Therefore, a cocktail of these two defined commensals is protective and can restore intestinal immunity to older animals.

## Discussion

From mammals to flies, aging is associated with dysfunction of the immune system, dysbiosis of the microbiota, increased inflammatory signaling, and elevated oxidative stress (Bischoff, 2016; Biteau et al., 2008; Buchon et al., 2009; Choi et al., 2008; Clark et al., 2015; Guo et al., 2014; Li and Jasper, 2016; Li et al., 2016; Man et al., 2014; Patrignani et al., 2014; Wong et al., 2011). Furthermore, aging increases susceptibility to pathogens including enteric viruses. The mechanisms that underlies these changes are poorly understood and are likely linked to the dysfunction of the immune system. While the microbiota is emerging as an important player in immunity and intestinal homeostasis, how changes in the microbiota impact immune function is only beginning to be understood. Moreover, how aging shapes immune functions via altered microbiota function is largely unexplored. This is in part due to limitations in manipulating the microbiota with age in vertebrate model systems. Flies have been used as a model for understanding the molecular mechanisms behind aging, which has revealed important insights into the aging of mammals (Bischoff, 2016; Li and Jasper, 2016; Man et al., 2014; Patrignani et al., 2014). Additionally, flies have emerged as a tractable system to study the microbiome (Broderick and Lemaitre, 2012; Charroux and Royet, 2012; Ma et al., 2015; Wong et al., 2011). By taking advantage of the genetic tools, manipulable microbiota and short lifespan, we have established a *Drosophila* model to explore age-dependent changes in intestinal immunity. We find that by middle age, flies become drastically more susceptible to diverse enteric viral infections. We cannot detect infection of a single enterocyte by VSV or DENV in the young gut, while we can readily detect infection in the aged gut (Figure 1A). This increased susceptibility is due, at least in part, to the aged microbiota, since loss of the microbiota decreases susceptibility (Figure 2A,B).

Indeed, a growing body of literature has shown that the intestinal microbiota impacts intestinal homeostasis, immune function, and susceptibility to viral infection. Importantly, we find that the microbiota is not a static force, but that depending on the composition of the microbiota, the signals provided, and the physiological state of the host, the microbiota can be protective or detrimental to enteric infection. We previously found that the microbiota confers protection to enteric viral infection in young flies, with Gram-negative commensals, through DAP-type peptidoglycan, priming immune function of enterocytes (Sansone et al., 2015). Additionally, this study demonstrates that the aged microbiota is detrimental to intestinal immunity. Without a better understanding of the signals provided by the constituents of the microbiota, it is difficult to interpret the role of the ‘generic’ microbiota in infection. Indeed, experiments in diverse systems have concluded that the microbiota confers protection (Cirimotich et al., 2011; Pang and Iwasaki, 2012; Ramirez et al., 2012; Schaffer et al., 1963; Varyukhina et al., 2012; Xi et al., 2008) or promotes (Carissimo et al., 2014; Jones et al., 2014; Kane et al., 2011; Karst, 2016; Kuss et al., 2011; Robinson et al., 2014) viral infection. Therefore, a better understanding of how particular constituents or consortia present within the microbiota shapes immunity in different contexts is essential to understanding what defines a healthy ‘microbiota’ and how we may intervene under conditions of dysbiosis to increase protective immunity.

Moreover, we found that we could revert the aging-dependent increase in susceptibility to enteric viral infection in older flies by reconstituting their microbiota with a healthy young flora using fecal transfers (Figure 3A). Characterization of the microbiota revealed that as flies age, there is increasing numbers of bacteria in their guts with a 100-fold increase observed at middle age (Figure S2A) and that there were striking changes in the relative compositions. Functional analysis of our metagenomic data revealed changes in ROS pathways, consistent with previous studies that found increased ROS with age (Buchon et al., 2009; Guo et al., 2014; Hochmuth et al., 2011). We characterized the ROS levels in the intestines of flies and found that the dysbiotic microbiota leads to increased ROS production: old or young flies harboring the microbiota from old animals had high ROS levels, while old or young animals with the microbiota from young animals did not (Figure 4D). Importantly, decreasing ROS production in older flies pharmacologically or genetically leads to a decrease in susceptibility to enteric viral infection (Figure 6A-C,E). Inducing high levels of ROS in young flies increased susceptibility to viral infection, suggesting that high ROS is sufficient to block immune function (Figure 5B). Collectively, these data demonstrate that ROS is a major rheostat for protective immunity in the intestine.

Our previous studies found that intestinal antiviral immunity was dependent on the induction of the antiviral cytokine Pvf2 (Sansone et al., 2015). Consistent with their increased susceptibility to infection, we found that older flies are unable to induce Pvf2 in response to viral challenge (Figure 1D, S1B). Moreover, forced expression of this cytokine in enterocytes of old animals was protective (Figure 1E). In addition, either by performing a fecal transfer with a young microbiota or by reducing ROS in older flies, Pvf2 expression is restored, leading to protective immunity (Figure 3D, 6F). Antiviral Pvf2 induction requires two signals: microbiota-dependent priming of NF-kB and virus-induced activation (Sansone et al., 2015). The aging *Drosophila* gut presents with increased inflammatory signaling (Figure S2B) (Buchon et al., 2009; Guo et al., 2014) consistent with observations in mammalian systems where aging is associated with inflammation, which has been termed inflammaging, and this hyperinflammatory state is likely contributing to the high levels of ROS (Franceschi et al., 2000; Frasca and Blomberg, 2016; Xia et al., 2016). Altogether, these data suggest that while the priming signal downstream of Gram-negative commensals remains intact in the aging intestine, high ROS is interfering with the virus-induced signal leading to increased susceptibility to enteric infection. Consistent with this, ROS depleted old flies that lack a microbiota were unable to induce Pvf2 upon infection (Figure 6F).

Since the young microbiota was protective, we set out to explore whether we could identify components of the microbiota that could confer protection to older animals. We monoassociated old animals with the two most prevalent Gram-negative or Gram-positive commensals. While the Gram-negative commensal *A. pomorum* was protective in young flies (Sansone et al., 2015), it was unable to provide protection in old flies (Figure 7A). In contrast, old flies reconstituted with the Gram-positive commensal *L. fructivorans* were less susceptible to viral infection than the conventional aged animals. While *L. fructivorans* was clearly protective, it did not restore complete immune function, as the flies remained more susceptible than conventional young animals (Figure 7B). DAP-type peptidoglycan present on Gram-negative commensals was required for Pvf2 induction in young animals (Sansone et al., 2015) and it is known that *L. fructivorans* does not have this type of peptidoglycan. Therefore, we reasoned that the lack of the peptidoglycan-derived signal was responsible for the difference in immune function between young conventional animals and old *L. fructivorans* monoassociated animals. Indeed, Pvf2 was not induced in older flies monoassociated with *L. fructivorans* (Figure 7C). Since heat-killed *A. pomorum* is sufficient to provide the priming signal for Pvf2 induction (Sansone et al., 2015), we tested whether gnotobiotic flies harboring *L. fructivorans* along with heat-killed *A. pomorum* were as resistant to enteric infection as young conventional animals and found that this combination was completely protective and restored cytokine expression (Figure 7B,C). Altogether these data suggest that monoassociation with *L. fructivorans* overcomes the ROS-induced block to immune function and that the combination of these two constituents can completely restore intestinal immunity to old flies.

How the aged or dysbiotic intestinal environment impacts enteric viral immunity in mammals is unknown. Moreover, how aging shapes immune functions via altered microbiota is largely unexplored. Our mechanistic studies provide insight into this biology and led to our discovery that the aging-dependent increase in susceptibility is not a terminal state. We can restore immune function in aged flies both pharmacologically and by altering the microbiota. Using pharmaceuticals or probiotics to promote a homeostatic and healthy gut environment will be possible once we have a clearer understanding of how the microbiota shapes immunity. Altogether, our studies demonstrate that the microbiome can be harnessed to alter the aging-dependent decline in enteric immunity.

## Materials and Methods

### Fly Rearing and Infections

All fly stocks used in this study are listed in Table S1. Flies were orally infected as previously described (Xu et al., 2013). In brief, female flies of the indicated age and genotype were orally infected with 10μL of each virus (DCV: 1x10^12^ IU/mL; VSV: 1x10^8^ pfu/mL; DENV-2: 2x10^8^ pfu/mL) in sucrose for three days and then transferred to virus-free food every 3 days or for the duration of the experiment. For survival, flies were scored daily for 20 days. Three independent replicates of 15 flies each were performed for each experiment. Heat shock flies were incubated at 37°C for 1 hour everyday for 3 days prior to infection. Once orally infected, flies were incubated at 37°C for 1 hour every other day for the duration of the experiment.

For drug treatment, flies were fed drug (DPI – 1 mM, Paraquat – 10mM) supplemented in 5% (vol/vol) in sucrose for three days on whatman. Flies were then infected as described above with the drug supplemented in 5% (vol/vol) in sucrose for the duration of the experiment.

For antibiotic experiments, female flies were transferred onto vials containing 200 μl of agarose-food (1.5% agarose, 7% corn syrup, 2% Bacto TC Yeastolate) supplemented with doxycycline (640 μg/ml), ampicillin (640 μg/ml), and kanamycin (1 mg/ml). Control flies were reared on agarose-food supplemented with vehicle. Three days later, flies were transferred to whatman paper containing 10 μl of virus for three days and transferred onto fresh antibiotic- or vehicle-containing food every 3 days for the duration of the experiment.

Defined commensals were grown in MRS broth at 29°C overnight (Newell and Douglas, 2014). Commensals from the feces of the age indicated flies were grown in MRS broth for 6 hours at 29°C. Fecal transferred and monoassociated flies were established by antibiotic treating and then transferring onto fly food amended with 5 x 10^8^ CFU of the indicated bacterial strains for 3 days. Oral infections were performed as described above. Commensals were heat killed by incubating at 80°C for 1 hour.

### Viruses, Chemicals, and Reagents

VSV-GFP, DCV, and DENV-2 were grown as described (Sessions et al., 2009; Xu et al., 2012). An antibody against DENV E protein (4G2) was provided by Michael Diamond (Washington University in St. Louis). Anti-DCV capsid antibody was used as described (Cherry and Perrimon, 2004). Additional chemicals were from Sigma.

### RNA and Quantitative Real-Time PCR

Total RNA was extracted from 15 fly guts, using TRIzol (Invitrogen) according to manufacturer’s protocol and as previously described (Xu et al., 2013). cDNA was prepared using M-MLV reverse transcriptase (Invitrogen). cDNA was analyzed using Power SYBR Green PCR Master Mix (Applied Biosystems), along with gene specific primers in triplicate, for at least three independent experiments. Data was analyzed by relative quantification, by normalizing to rp49. Primers are listed in Table S2.

### Shotgun Metagenomic Sequencing and Analysis

DNA was extracted from 8 x 10^8^ CFU bacteria from fly feces using PowerSoil DNA Isolation Kit (Mo Bio Laboratories) according to manufacturer’s protocol. The genomic DNA was processed for shotgun metagenomic sequencing using the NexteraXT Library Preparation Kit (Illumina) according to manufacturer’s recommendations. The library was sequenced using single-ended on a NextSeq platform (Illumina) with a read length of 150bp. Sequence reads were quality-trimmed using Timmomatic (Bolger et al., 2014). CosmosID databases (Rockville, MD) were used to assign bacterial taxonomy and principal component analysis to the metagenomic reads. Trimmed reads were fed through the Functional Mapping and Analysis Pipeline (FMAP, PMID: 27724866) (Kim et al., 2016) for alignment, gene family abundance calculations, and statistical analysis, using the default UniRef100 gene and KEGG orthology database. RPKM values are of pathways of interested were manually tallied.

### Immunofluorescence

Guts were processed as previously described (Xu et al., 2013). Briefly, 5 guts per experiment were dissected in PBS, fixed in 4% formaldehyde solution for 30 minutes, rinsed 3 times in PBS, and blocked with 5% normal donkey serum for 45 minutes. Samples were incubated with primary antibody (DCV capsid 1:3000) or (Dengue 1:4000) overnight at 4^°^C, rinsed 3 times in PBT, and incubated with secondary antibody (1:1000) and Hoescht 33342 (1:1000) at room temperature for 1 hour 15 minutes. Samples were rinsed 3 times in PBT and mounted in Vectashield (Vector Laboratories). Guts were imaged on Leica TCS SPE confocal microscope at 40X. Three independent experiments were performed imaging.

### ROS analysis

Flies were overnight starved on whatman paper with a 1:1 mixture of water and PBS. The following morning flies were starved for 1 hour in empty vials to synchronize feeding. Flies were feed 100 μM ROSStar 550 (LI-COR Biosciences) for 4 hours. Five guts were dissected in cold PBS, fixed in 4% formaldehyde solution for 5 minutes, rinsed 3 times in PBS, incubated in PBT with Hoescht 33342 (1:1000) for 5 minutes, rinsed 3 times in PBS, and mounted in Vectashield (Vector Laboratories). Guts were imaged on Leica TCS SPE confocal microscope at 40X. Three independent experiments were performed.

### Bacterial Plating

Five guts per sample were dissected under sterile conditions in PBS and crushed. Serial dilutions were plated on MRS agar plates and incubated at 29^°^C for two days. Three independent experiments were performed.

### X-Gal Staining

X-gal staining was performed as previously described (Choi et al., 2008). In brief, 5 guts per experiment were dissected in PBS and fixed in 1% glutaraldehyde for 10 minutes. Samples were washed 3 times in PBS and stained with 0.2% X-gal in staining buffer (6.1 mM K_4_Fe(CN)_6_, 6.1 mM K_3_Fe(CN)_6_, 1 mM MgCl_2_, 150 mM NaCl, 10 mM Na_2_HPO_4_, 10 mM NaH_2_PO_4_) in the dark at room temperature.

### Statistics and Data Analysis

For survival curves, pair-wise comparisons of each experimental group with its control were carried out using a Mantel-Haenszel test. For RT-qPCR studies, P values were obtained by comparing delta CT values for three independent experiments except Figure 7C where a two-way ANOVA was performed. For other experiments, the Student’s two-tailed t-test was used to measure the statistical significance in each experiment and then considered significant if p<0.05.

## Acknowledgements

We would like to thank the following for fly stocks: Bloomington, Kyoto, W.J. Lee, N. Perrimon, and M. Yoo. We would like to thank M. Diamond for antibodies; J. Rose, R. Hardy, P. Christian, and ATCC for viruses. This work was supported by National Institute of Health grants 5T32AI055400 to C.S. and RO1AI122749, RO1AI095500, and RO1AI074951 to S.C. and pilot funding from P30DK050306. S.C. is a recipient of the Burroughs Wellcome Investigators in the Pathogenesis of Infectious Disease Award.

